# Deformable mirror-based two-photon microscopy for axial mammalian brain imaging

**DOI:** 10.1101/736124

**Authors:** Alba Peinado, Eduardo Bendek, Sae Yokoyama, Kira E. Poskanzer

## Abstract

This work presents the design and implementation of an enhanced version of a traditional two-photon (2P) microscope with the addition of high-speed axial scanning for live mammalian brain imaging. Our implementation utilizes a deformable mirror (DM) that can rapidly apply different defocus shapes to manipulate the laser beam divergence and consequently control the axial position of the beam focus in the sample. We provide a mathematical model describing the DM curvature, then experimentally characterize the radius of curvature as well as the Zernike terms of the DM surface for a given set of defocuses. A description of the optical setup of the 2P microscope is detailed. We conduct a thorough calibration of the system, determining the point spread function, the total scanning range, the axial step size, and the intensity curvature as a function of depth. Finally, the instrument is used for imaging different neurobiological samples, including fixed brain slices and *in vivo* mouse cerebral cortex.

## 1. Introduction

Two-photon (2P) microscopy[1,2] is a widely used technique for imaging in high scattering media, such as the mammalian brain. This technique allows imaging in deep brain tissue, taking advantage of low scattering and absorption in the near infrared range. Much current 2P microscopy takes advantage of genetically encoded calcium (Ca^2+^) indicators to record neural activity in the brains of head-fixed animals[3]. While most 2P calcium imaging targets neuronal activity, other cell types also exhibit spontaneous Ca^2+^ dynamics. These include astrocytes, the largest class of non-neuronal brain cells which are largely electrically silent but which exhibit spatiotemporally rich Ca^2+^ dynamics[4].

2P microscopy is a laser scanning technique in which a *xy* scanner is used to move the beam focus across a sample, obtaining a 2D image. Recently, a variety of volumetric 2P systems that enable population-wide imaging at multiple *z*-planes have been reported to enable the study of neural activity across brain tissue depths[5]. One group of these systems applies technologies that sequentially move the focus along the *z* axis, and includes piezo-actuated objectives[6–8], electro-tunable lenses[9–11] and remote focusing[12–14]. Because their settling times are on the order of a few milliseconds, these technologies are not fast enough to capture relevant *in vivo* depth-imaging with high *z*-resolution, limiting their use to in neural physiological imaging only a few defined *z* planes. Other systems use a high-speed axial scanner such as an acoustic-optical scanner[15]or an ultrasound-driven lens[16], but they can suffer from focusing aberrations. Another solution to imaging in depth is the technique of implanting microprisms in the brain to image orthogonal to the skull surface while still using a traditional *xy* scanner[17,18]. For cell types, including astrocytes, which become reactive upon injury, this type of invasive technique may not allow the study of physiological—rather than pathological—dynamics. Still other techniques for 2P imaging beyond two dimensions are based on spatially multiplexing the beam focus at different depths to carry out volumetric 2P *in vivo* brain imaging. Some use a Spatial Light Modulator (SLM) to split one beam into several beamlets at different depths [19], while others use Bessel beams[20,21] to simultaneously excite an axially elongated focus. However, the recorded signal for these instruments is a 2D projection of a 3D volume. Therefore, they depend on source-separation algorithms[22] to retrieve individual signal contributions, built expressly for neuronal somatic activity. However, astrocytic Ca^2+^ activity exhibits differential characteristics from neuronal Ca^2+^ activity with more complex and variable spatial dynamics, and these characteristics render current source-separation algorithms unusable for astrocytes. For all these reasons, new imaging techniques are necessary to record activity of neuronal and non-neuronal cell types across layers with a high *z*-resolution.

Here, we use a deformable mirror (DM) as an active optical element to achieve fast axial scanning for *in vivo* brain imaging. A DM has a surface that can be dynamically shaped in order to control the light wavefront. Recently, MEMS (micro-electro-mechanical systems)-based DMs have been shown to be capable of changing their surface shape at KHz rates[23]. Using this technology in a 2P microscope allows us to move the laser focus along the *z* axis, beating current *z*-scanning technologies. Moreover, because the DM is a reflective optical element, the optical setup is achromatic and can be applied for the whole broadband pulsed laser spectrum. In this paper, we implement a MEMS DM-based 2P microscope for achieving high-speed axial scanning in mammalian brain in order to study any Ca^2+^-signaling brain cell, including neurons and astrocytes. This work complements the contributions of others in the field [24–26]. For example, a 2P microscope with axial control[25] has been developed, but not applied to image biological samples. Also, a DM-based 2P microscope has been implemented for volumetric imaging neurons in *Drosophila* [26], but not for imaging the central nervous system of mammals, including neurons and non-neuronal cells.

The outline of this paper is as follows: In Section 2, we introduce the concept of using the DM as a fast axial scanner, present a model for its use, and experimentally calibrate its performance. In Section 3, the optical setup of the DM-based 2P microscope is described and an experimental characterization is provided. In Section 4, experimental results of the instrument are given, including imaging of pollen grains, fixed brain slices, and *in vivo* mouse cerebral cortex. Finally, the last section summarizes the main results of this work.

## 2. Deformable mirror technology for fast axial scanning

### 2.1. Concept

A DM is a reflective device that can modify the phase of an incident electromagnetic wavefront by changing the shape of its surface. An array of actuators adjusts the axial position (*z*) at their given location on the DM surface plane (*x,y*). The intrinsic reflective operation of the device results in a wavefront change twice as large as the surface stroke applied. There are segmented DMs (composed of independent flat mirror segments) and continuous DMs (with actuators fixed to the back side of a continuous reflective membrane). Nowadays, many technologies use DMs as the key element for applying active and adaptive optics in diverse applications, such as astronomy[27], ophthalmology[28], and biology[29,30]. In applications requiring smooth wavefront control, a continuous DM is advised. Fig. 1(a) shows a diagram of this type of continuous DM.

**Fig. 1.**
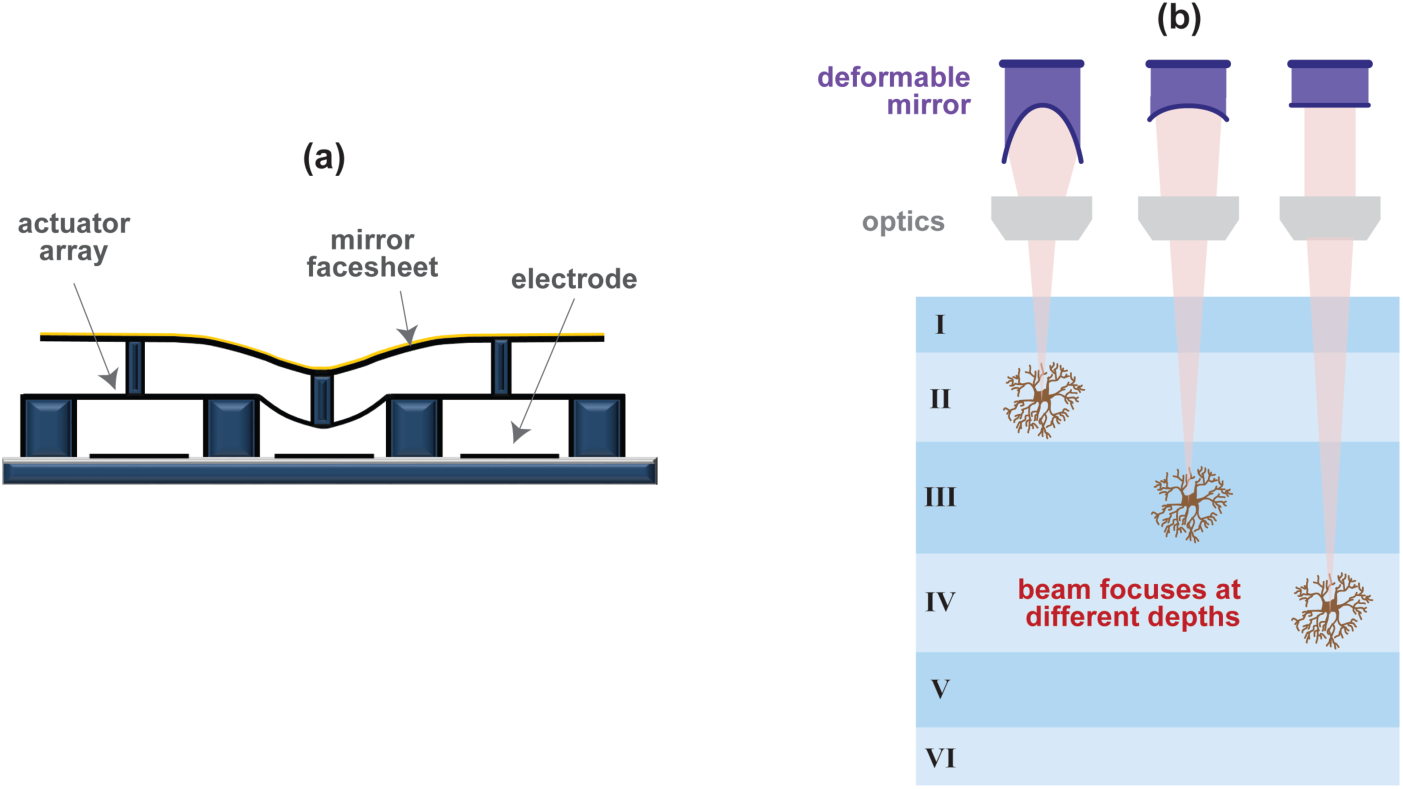
(a) Continuous DM structure: electrodes are used to move an array of actuators, which are covered with a mirror facesheet. (b) Concept sketch: A DM is used as a dynamic element for fast axial scanning during brain imaging, allowing imaging of different cortical layers.

In this work, we develop a method for recording brain activity at different neural depths in which a DM is used to change the incoming wavefront to the microscope objective. This allows axial movement of the focal point along the sample, and results in 2P imaging at different depths. Together, this leads to a novel fast axial scanning, DM-based method for multi-layer brain imaging, a concept schematized in Fig. 1(b).

### 2.2. Model

A lens modifies the phase of an incoming wavefront when light is transmitted through it. If the incoming beam is collimated, the lens will change the wavefront phase in such way that at its exit pupil, there will be a spherical wavefront converging to a point a distance *R*, where *R* is the focal length of this lens.

When a defocus term,

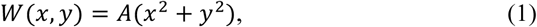

is added to the original spherical wavefront of radius of curvature *R*, a focal shift will result given by[31]:

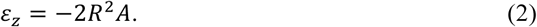

*A* is a parameter that defines the sign and amplitude of the defocus, and consequently it also modifies the axial focal shift. Therefore, when the optical path difference (OPD)[31] at the entrance pupil increases because of the addition of a positive defocus term (*A* > 0), the spherical beam converges to a focus point moved toward the lens in the negative *z* axis direction. In contrast, for a negative defocus term (*A* < 0), the focus point is moved away from the lens in the positive *z* axis direction.

To apply the same wavefront change to the entire field-of-view imaged through the system, the DM must be placed at a plane conjugated to the microscope objective entrance pupil. In that way, when a quadratic surface is implemented on the DM, the reflected wavefront has a defocus term, which is transferred to the entrance pupil of the microscope objective causing a controlled axial shift of the focal point.

We implemented a set of defocus distributions using Eq. (1), and we changed parameter *A* from −1 to 1, corresponding to mirror shapes from the maximum negative to positive radius of curvature. In particular, when *A* = −1 (+1) the DM has a concave (convex) shape of radius of curvature −1.14 m (+1.14 m). The surfaces were modeled with respect to a flat surface in which all actuators were at half of their maximum displacement (stroke) remaining after flattening the DM. Each DM requires a specific voltage map to flatten its surface.

In summary, the process of obtaining the final voltage map to be sent to the DM is the following (Fig. 2): First, a set of ideal defocuses, *z(x,y)*, are modeled using Eq. (1) with a quadratic phase and a parameter *A* which defines the sign and magnitude of the radius of curvature. These functions, defined in the deflection space, are added to the flat map, *z*_*flat*_*(x,y)*, and as a result, the corrected defocus deflection distribution, *z’(x,y)*, is obtained. Then, these deflection distributions are converted from the deflection space to the voltage space by using the deflection curve, a calibration curve provided by the DM manufacturer which relates voltage applied to a DM actuator with the actual deflection. Finally, we obtained the corrected defocus distribution in voltage units, also referred as voltage map.

**Fig. 2.**
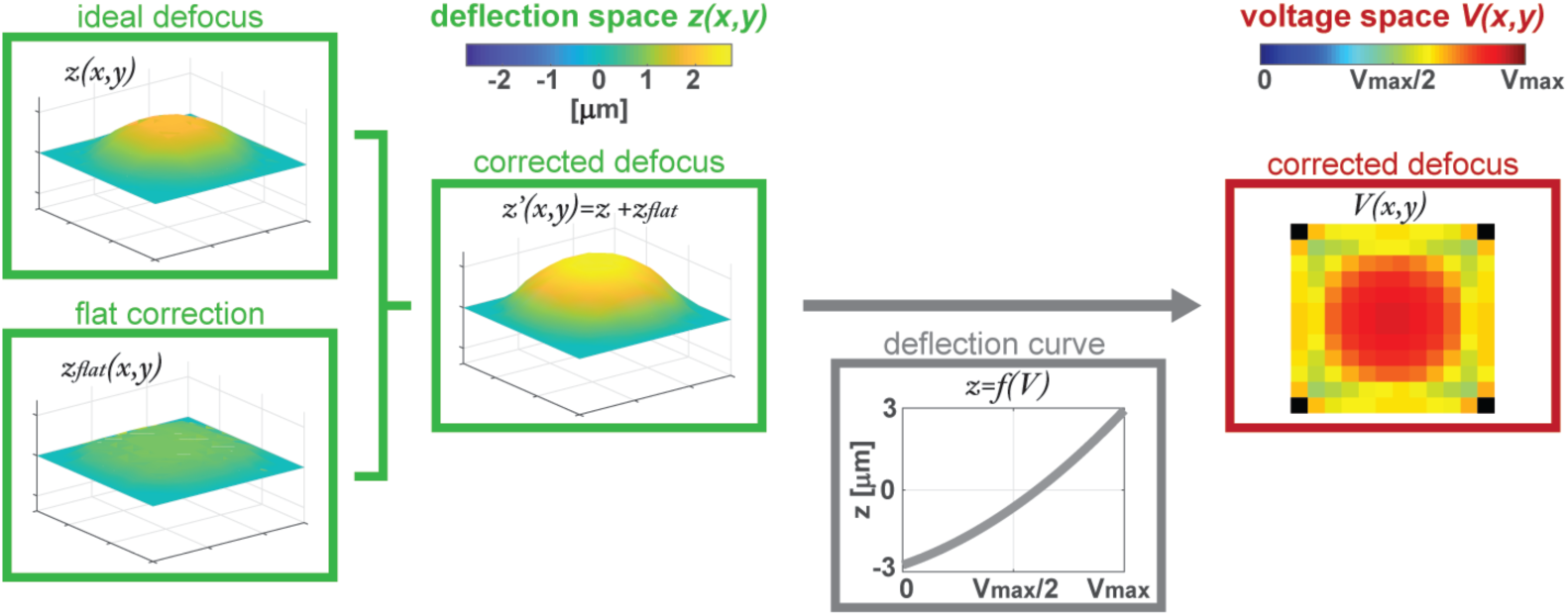
Flow chart describing the process from the ideal defocus deflection distribution to obtain the voltage map sent to the DM, in which we consider the flat correction needed to flatten the intrinsic DM bow and the DM deflection curve.

The radius of curvature (*R*) of the DM defocus shape can be expressed as a function of stroke (*s*), and DM semi-aperture (*x*), using the intersecting chord theorem:

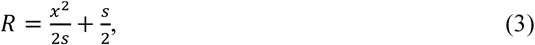

where the stroke is the peak-to-valley from the ideal defocus deflection distribution.

A variety of DMs are commercially available, and these vary by size, continuous versus segmented type, actuator pitch and stroke, number of actuators, and response time, with a trade-off between stroke and response time. For this project, we selected a continuous DM (Multi-5.5 DM, Boston Micromachines Corporation) of 12×12 actuators matrix with a maximum stroke of 5.5 µm and a total aperture of 4.95 mm, which offers a compromise between stroke and response time, which for this DM is < 100 µs. The four actuators at the DM corners are fixed, for a total of 140 operative actuators. In addition, we used a circular mask of radius 5.5 actuators in which all actuators inside that mask are defined as the values given by protocol from Fig. 2, whereas actuators outside of it are assigned to the flat map value. This mask helps to ensure the implementation of the desired shape inside the pupil even though the bordering actuators are fixed.

### 2.2 Experimental characterization

#### 2.3.1. Free propagation

In this section, we describe an experimental characterization of the focusing capability of the DM when implementing a concave mirror (*R* < 0) with the DM. To this aim, we implemented the following set-up (see Fig. 3(a)): A collimated beam is used to illuminate the DM, and the reflected beam is propagated. Then, we measured the distance in which the beam was focused from the DM. In other words, we measured the experimental focal length, *f*_*exp*_, of the DM acting as a converging lens. From the focal length measurements, we calculated the radius of curvature:

**Fig. 3.**
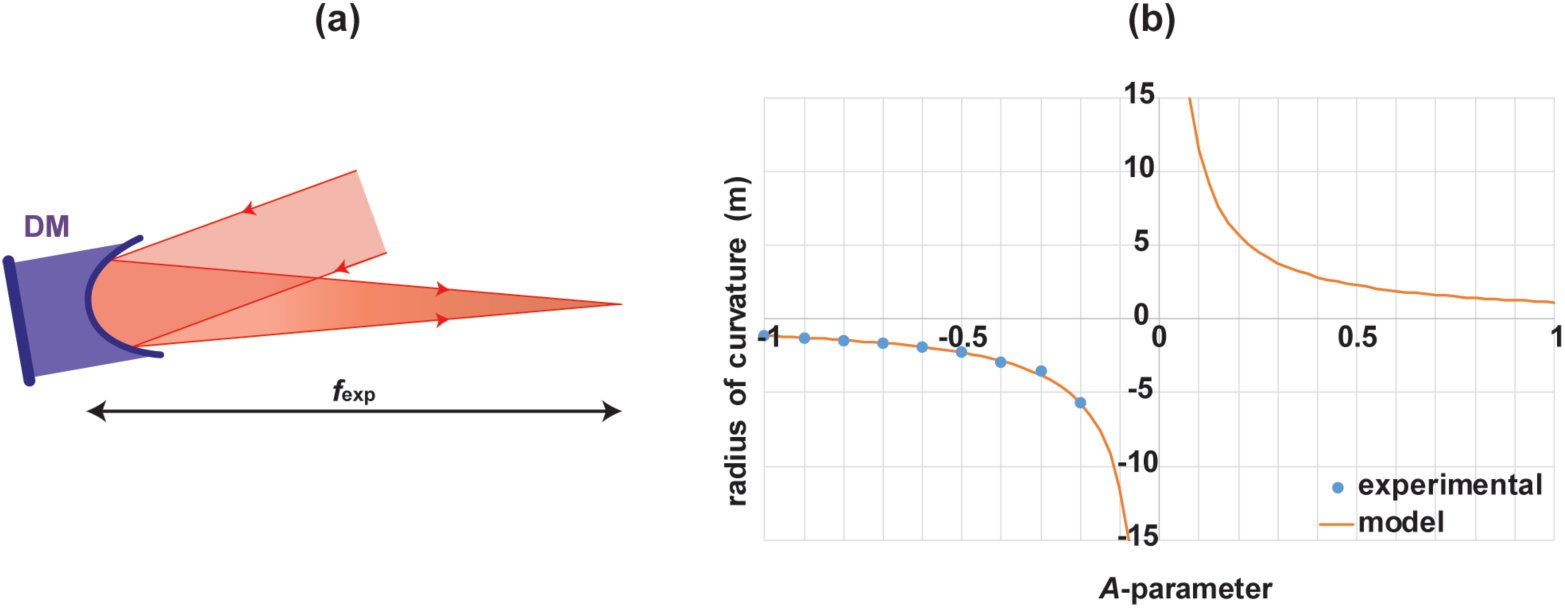
(a) A collimated light beam is focused at a distance *f*_*exp*_ by the DM with a concave shape. (b) Experimental and theoretical radius of curvature of the DM as a function of the *A*-parameter.

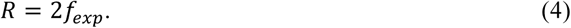

DM radius of curvature as a function of the *A*-parameter is plotted in Fig. 3(b). Blue points correspond to the experimental values using Eq. (4). In addition, the continuous orange line corresponds to the modeled data using Eq. (3). When *A* < 0, radius of curvature is negative too, corresponding to a concave mirror; then the reflected beam converges at some point and *f*_*exp*_ could be measured. Whereas when *A* > 0, radius of curvature is positive too, corresponding to a convex mirror, and the light beam gets divergent after the DM reflection and it does not get focused, so no experimental points were acquired. Finally, when *A* = 0, corresponding to a flat DM, the radius of curvature is infinite. Thus, the DM does not affect the convergence of the reflected beam. We observe a good agreement between the experimental data and model, proving that the DM manipulates the wavefront as expected when using converging-like curvatures. Experiments were conducted illuminating with a red diode laser at 635 nm (CPS635R, Thorlabs) but the DM is achromatic, so the experiment is relevant on the infrared as well.

#### 2.3.2. Wavefront calibration

After we validated the focusing DM capability for positive convergent-like shapes, we tested arbitrary DM defocus shapes by measuring the actual reflected beam wavefront with a Shack-Hartmann wavefront sensor[32]. Any wavefront can defined by a linear combination of the Zernike polynomials[31], a basis of optical aberrations whose terms are orthogonal over the interior of a unit circle.

The experiment consisted of illuminating the DM with a collimated laser beam at 635 nm (CPS635R, Thorlabs), and immediately measuring the reflected beam wavefront using a Shack-Hartmann wavefront sensor (WFS30-5C, Thorlabs). 21 different shapes were implemented with the DM, varying the *A*-parameter from −1 to 1 in steps of 0.1. The wavefront was measured as a function of the Zernike polynomial terms. The weights measured for the different Zernike coefficients are plotted in Fig. 4 for the 21 DM shapes (given by *A*-parameter). To help the reader visualize each Zernike term, Fig. 4 also includes in the horizontal axis the wavefront map of each Zernike term in a unitary circle. The defocus term, highlighted with a pink box in Fig. 4, is linearly dependent with the *A*-parameter, whereas the other Zernike terms are negligible.

**Fig. 4.**
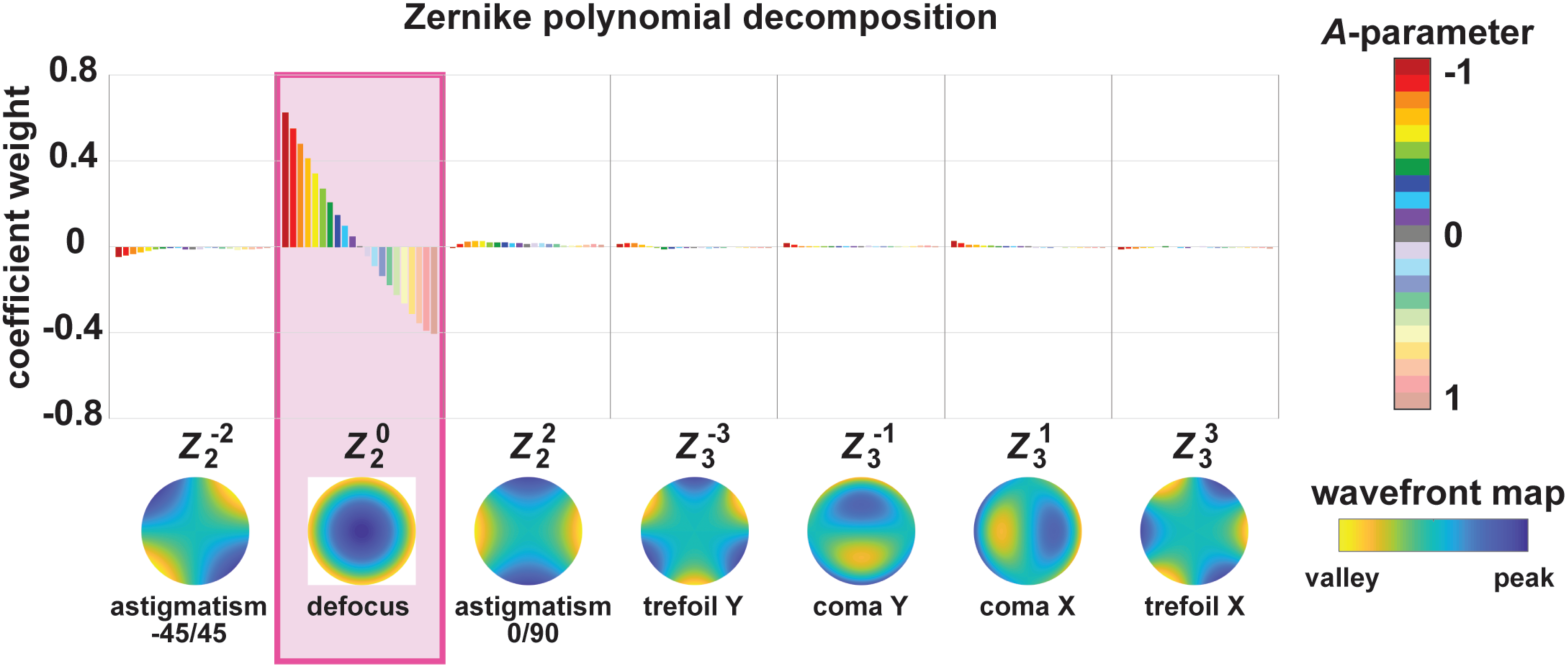
Zernike polynomial decomposition for 21 different DM shapes (*A*-parameter) measured with a Shack-Hartmann wavefront sensor.

## 3. Experimental setup

### 3.1. Optical layout

Our 2P microscope is a custom-built instrument which combines a traditional upright fluorescence microscope (BX51WI, Olympus) with three additional modules to enable 2P microscopy: a laser source including the relay optics, an active optics module where the DM and scanning galvanometer mirror set are housed, and a detection system that uses Photomultiplier Tubes (PMTs).

The components are shown in Fig. 5(a). First, a Ti:Sapphire laser (Mai Tai HP, Spectra Physics) is used as an illumination source, with a tunable wavelength from 690 to 1040 nm and pulses of < 100 fs with a pulse rate of 80 MHz. In addition, a Pockels cell (350–80, Conoptics) controls the beam intensity illuminating the sample. A beam relay system delivers the laser beam at a reimaged pupil located at the surface of the DM. The DM is used to implement different curvatures to manipulate the wavefront (Fig. 5(b)), and consequently, to generate different displacements of the beam focus along the *z* axis beyond the microscope objective (Fig. 5(c)). Then, *xy*-scanning galvanometer mirrors are used (6215H, Cambridge Technology) to scan the focal point over the *xy* plane on the biological sample. A set of lenses are used to reimage the DM surface to a pupil located between the galvanometer mirrors. A third, and last, reimaged pupil is formed using another set of lenses on the entrance pupil of the objective. All lenses, distributed by Thorlabs, are achromatic in the range of 650–1050 nm. Their focal lengths were selected to ensure the proper magnification at each reimaging process. All mirrors used are achromatic in the NIR range, optimized for ultrashort pulsed lasers (10B20EAG.1, Newport). A dichroic mirror is placed at 45 degrees above the microscope objective to transmit wavelengths > 660 nm, allowing light transmission from the IR laser to the sample, but reflecting emitted light in the visible range (as a result of the 2P process) from the sample to the detector module. The detector module is composed of, first, an IR blocking filter (ET680sp-2p8, Chroma) followed by a single-band bandpass filter (Semrock) and finally, a PMT (R6357, Hamamatsu). Finally, our system is controlled using ScanImage[33], an open-source MATLAB-based software for laser scanning microscopy focused on neurobiology. The microscope has the flexibility to be operated with different microscope objectives, but experimental data shown in this paper was obtained using a 20x water-immersion objective with NA 1.0 (XLUMPlanFLN, Olympus).

**Fig. 5.**
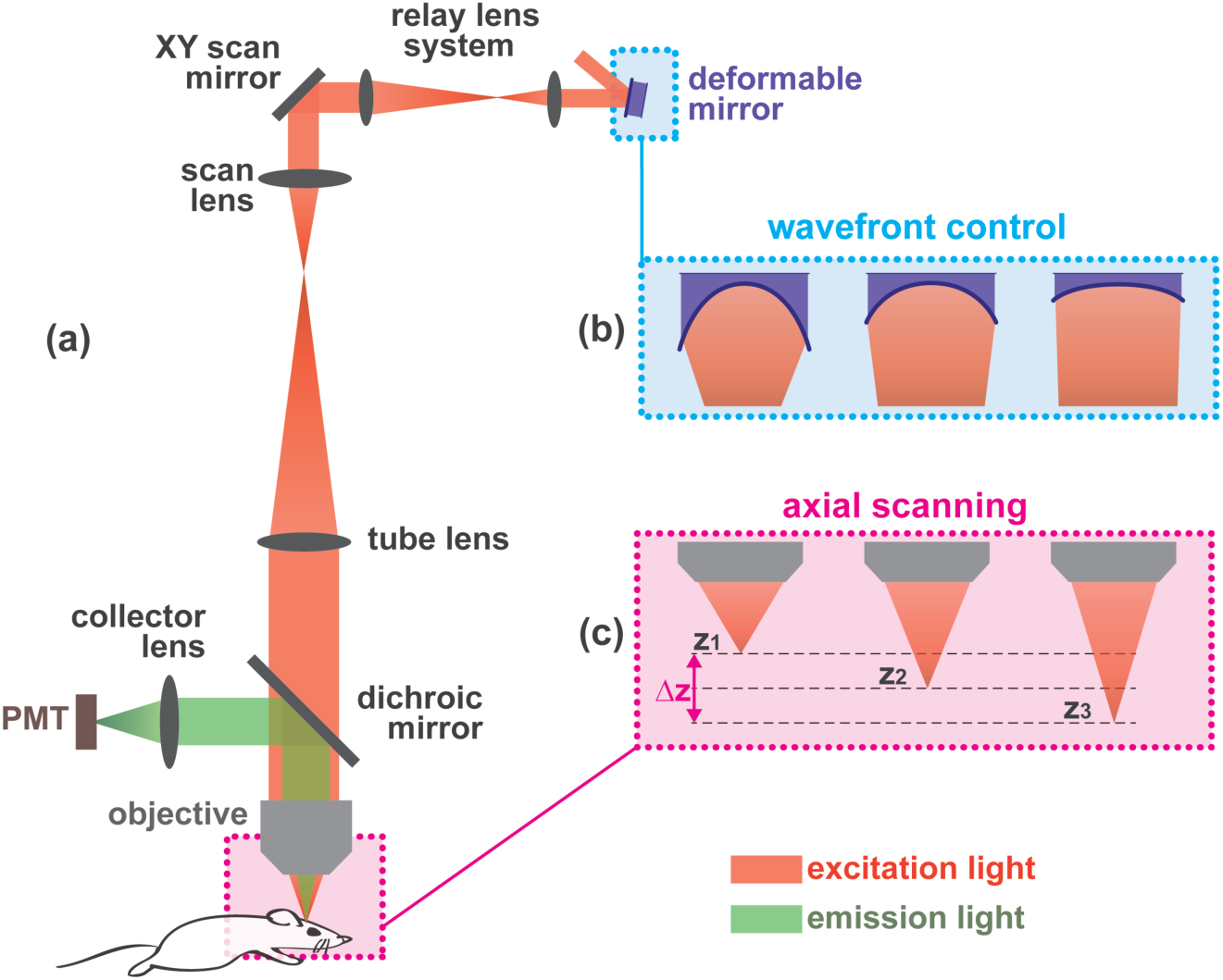
(a) 2P microscope optical setup. Close detail of the DM-based *z*-scanner, in which (b) a DM is used for manipulating the light beam wavefront, which causes (c) different axial displacements of the beam focus over the specimen.

The optical layout has been modeled in ZEMAX, to establish the baseline design that includes pupil relays and the mirror scanning system. The ZEMAX optimization tool was utilized to fold the optical system in the space available and using off-the-shelf components, obtain the component parameters such as the focal lengths, diameters, and distances of the lenses. In addition, the ZEMAX model was used to predict the performance of the instrument. We determined the theoretical *z* position of the beam focus for different curvatures of the DM for our given optical layout. In so doing, we estimated that the total axial scanning range (*Δz*) is 63 µm, achieved when applying the two maximum and opposite curvatures of the DM. Note that *Δz* could be increased by using a smaller NA for our system, but compromising the lateral spatial resolution. We also explored the impact of depth-scanning on the optical train. For example, when the focus is moved deeper into the sample, the laser beam at the entrance pupil of the objective is diverging. Thus, it has a larger diameter that causes vignetting at the edge of the aperture, which is one of the limiting factors of the scanning depth.

### 3.2. Experimental characterization

The DM-based 2P microscope has been implemented as described above and experimentally characterized. First, an analysis of the point spread function (PSF) as a function of different DM curvatures is detailed. In particular, we characterized the lateral and axial spatial resolution. Next, we determined the *z* scanning capability with the DM in terms of total scanning range and step size. Finally, the dependence of the intensity reaching the sample as function of *z* is studied.

#### 3.2.1. Point spread function

The PSF of the instrument was calibrated by imaging fluorescent microspheres (F8803, Invitrogen). These 0.1 µm-diameter beads have one-photon excitation and emission wavelengths at 505 and 515 nm, respectively. A solution of 4% low melting point agarose and water was used to dissolve 0.5% of the microspheres and then this was solidified on a glass slide and then coverslipped. The microsphere sample was illuminated with the pulsed laser at 1020 nm, and a long-pass filter at 660 nm was used to transmit that light to the sample and reflect to the detector the reemitted light after 2P absorption.

The protocol applied to characterize the PSF of our instrument was the following: For each DM curvature, a *z*-stack of 160 *xy* images was acquired while moving the microscope objective in steps of 0.25 µm along the *z* axis using a linear motorized stage (DRV014, Thorlabs).

As an example, we show in Fig. 6(a), (b), and (c) the PSF cross sections corresponding to planes *xy, yz* and *xz*, respectively, when the DM is in the flat configuration, so no curvature is applied (*A* = 0). The procedure is repeated for 4 different DM curvatures (*A* = −0.5, −0.25, 0.25 and 0.5). Experimental data was processed using MetroloJ [34,35], an ImageJ plugin to characterize the instrument’s PSF. The algorithm fits all three profiles (corresponding to the *x, y* and *z* axis) to a Gaussian function and determines the resolution in terms of Full Width at Half Maximum (FWHM) for each axis, based on the fitting parameters. Axial and lateral resolutions were characterized for all 5 analyzed DM curvatures, and values are given in Table 1. Results show that axial resolution is larger than lateral resolution, as it is well-known for 2P microscopy [2,5]. Moreover, Table 1 shows that axial resolution is more sensitive to a change in the DM curvature compared to the lateral resolution, which remains similar. Note that as the *A*-parameter gets closer to −1, the reflected beam converges and its diameter at the microscope objective entrance pupil decreases. Because the microscope objective is underfilled, the effective NA decreases, causing an increase in the axial resolution[2], a dependency shown in Table 1.

**Fig. 6.**
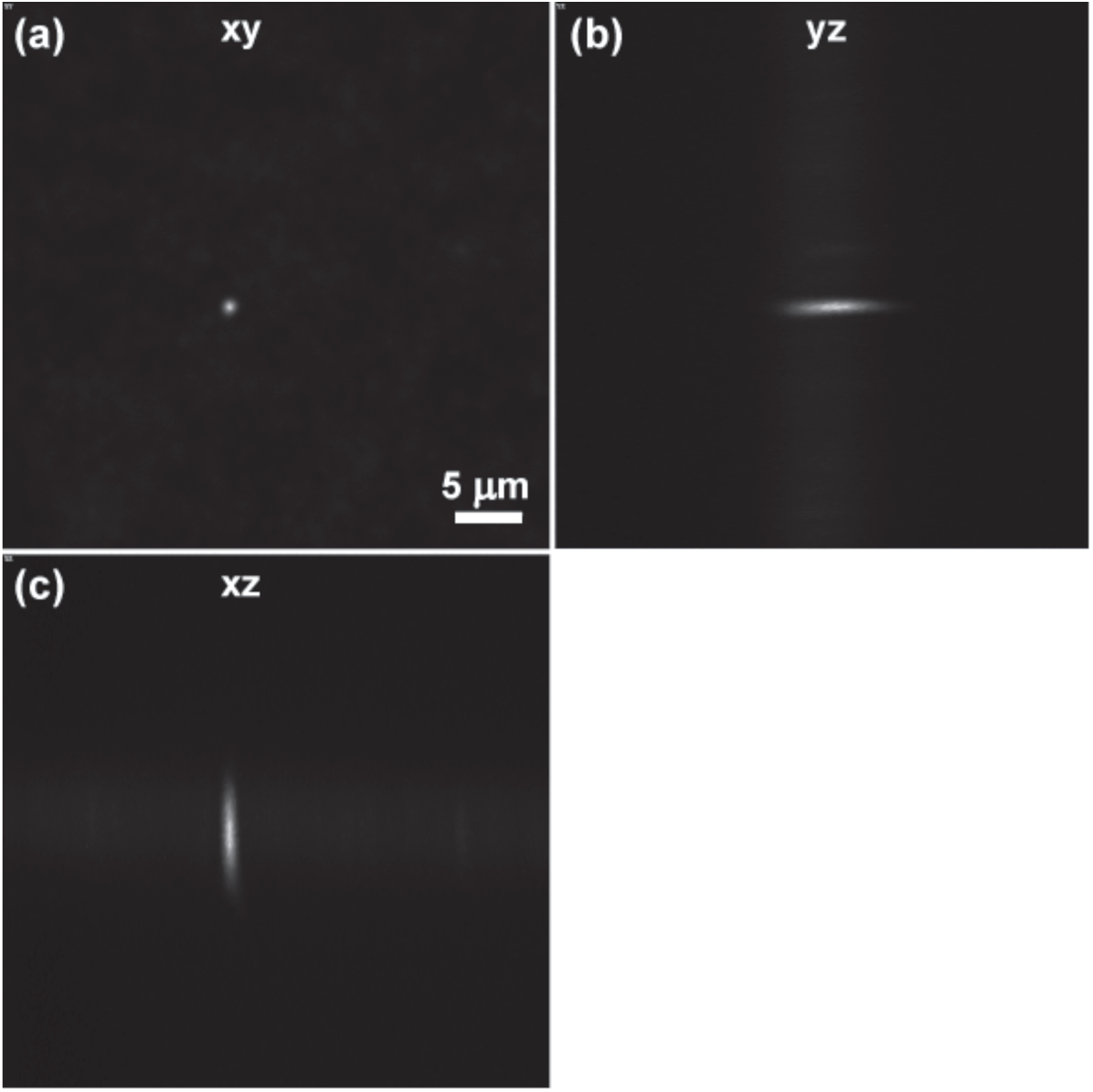
Experimental point spread function (PSF) when measuring 0.1 µm-diameter microspheres at 1020 nm, when the DM is flat (*A* = 0). (a) *xy*, (b) *yz*, and (c) *xz* planes.

**Table 1.** Lateral and axial spatial resolution experimentally measured for different curvatures used with the DM and with an illumination of 1020 nm. Spatial resolutions given as FWHM. *A*-parameter determines the DM curvature.

| $A$-parameter | - 0.5 | - 0.25 | 0 | + 0.25 | + 0.5 |
| --- | --- | --- | --- | --- | --- |
| FWHM x ($\mu\text{m}$) | 0.86 | 0.79 | 0.88 | 0.93 | 0.92 |
| FWHM y ($\mu\text{m}$) | 0.89 | 0.93 | 0.91 | 0.88 | 0.89 |
| FWHM z ($\mu\text{m}$) | 6.39 | 5.85 | 5.77 | 5.42 | 5.27 |

#### 3.2.2. Axial scanning capability

With the same experimental data, we next determined the *z* coordinate in which the PSF is centered by plotting the PSF *z*-profile in Fig. 7(a) (experimental data in circles and gaussian fit as a continuous line), and found the *z*-position of the peak. The peak *z*-coordinate values are plotted in Fig. 7(b). In addition, we added a linear regression fit equation, which relates the *z* position as function of the *A*-parameter. In other words, it describes how the DM is axially scanning in depth along the sample as a function of the DM curvature, given by the *A-* parameter, and this relation is:

**Fig. 7.**
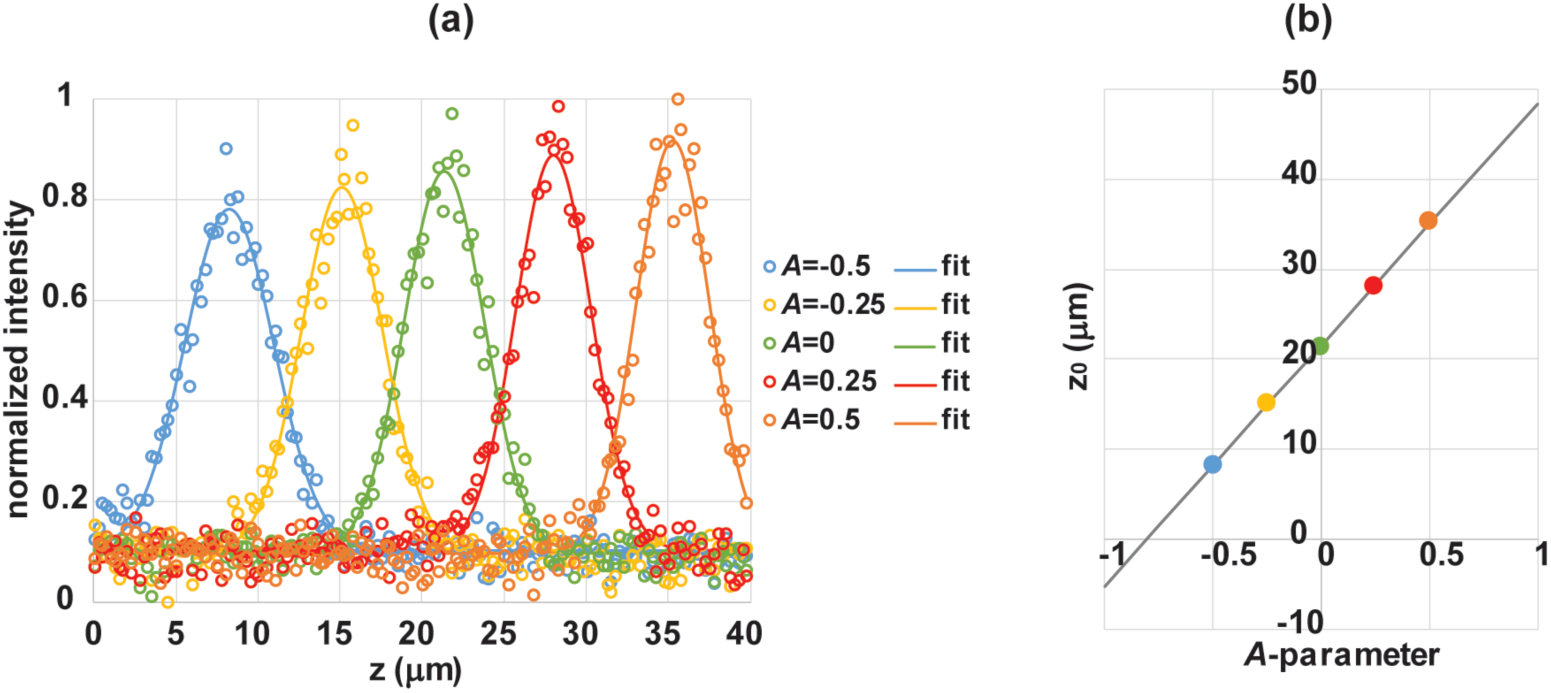
(a) Cross-section of the PSF along the *z* axis for different DM curvatures: experimental data (circles) and gaussian fit (continuous lines). (b) Intensity *z*-profile peak (*zo*) as a function of the *A*-parameter. Color points correspond to experimental data and continuous line is a linear fit (linear fit equation given in Eq. (5), with a R^2^ = 0.9994).

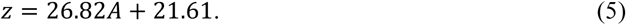

Using Eq. (5), we determined that the total experimental scanning range of our system (*Δz*) is 54 µm, obtained from the *z* difference between *A* equal −1 and +1. This is slightly smaller than the total scanning range value calculated with the optical model, because the DM needs an initial voltage map to be applied to compensate an initial bow when unpowered, leading to a slight decrease of the effective stroke for some of the DM actuators.

The axial step size (*δz*) that we can achieve with the DM scanner will depend on the *A*-parameter step size (*δA*) used for defining the DM curvature maps. In our experimental measurements, we used two sets of DM curvatures with different *δA*. First, a set of 21 curvatures was used in which δ*z* was 2.7 µm. Later, in order to have a finer depth resolution, we worked with a set of 201 DM curvatures with a δ*z* of 0.3 µm. Note that this axial scanning step could be even smaller if needed, only requiring a new set of DM curvatures from a smaller *δA.*

#### 3.2.3. Intensity curvatures

Finally, we observed a slight decrease in intensity when imaging at deeper layers. To understand and quantify the intensity lost with increasing *z*, we conducted the following test: we placed a uniform fluorescent slide under the objective and imaged at different depths. Two methods were used for axial scanning. One, using the DM with different curvatures (*δA = 0.1, δz = 2.7* µm), and two, the DM was kept in its flat configuration (*A* = 0) and a linear stage was used to axially move the microscope objective in steps of 2.5 µm. The pixel intensity mean of the acquired images as a function of *z* is plotted for both methods in Fig. 8. There is an intensity decrease when imaging at deeper layers using the DM compared to the linear stage. When we are imaging at deep layers (*A* > 0), the beam reaching the objective is divergent. Consequently, the microscope objective entrance pupil is overfilled, vignetting occurs, and only a fraction of the beam is transmitted through the objective lens to the sample, causing the reduction in intensity observed in Fig. 8 at deeper layers (large *z*). This intensity loss as *z* increases can be compensated by increasing the incident beam power using the Pockels cell. In fact, using the calibration curve plotted in Fig. 8, one could compensate in such a way that the intensity reaching the sample is kept constant, independently of the curvature of the DM.

**Fig. 8.**
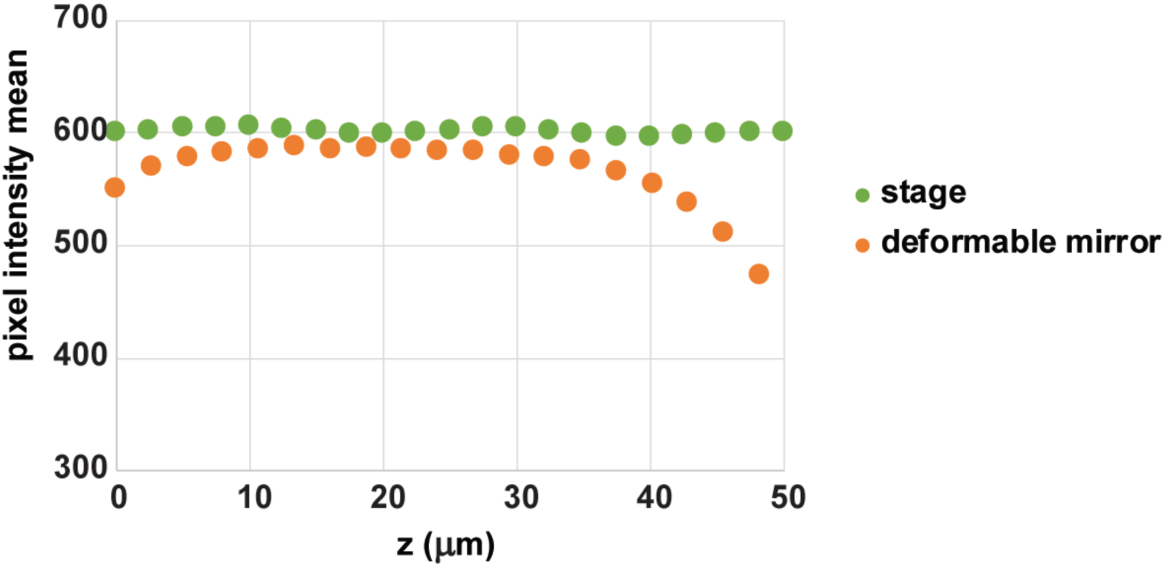
Pixel intensity mean, when imaging a uniform fluorescent slide at 950 nm, as a function of *z*. Depth is changed by moving the objective with a stage (green points) or with the DM applying different curvatures (orange points). Increasing *z* corresponds to image a deeper layer (*A*-parameter from the DM gets closer to 1).

## 4. Experimental measurements

The instrument has been tested with three different samples: pollen grains, a fixed mouse brain slice, and lastly *in vivo* mouse cerebral cortex.

### 4.1. Pollen

The sample was composed of grains of different types of pollen. 4 images were acquired at different depths, 8 µm apart. The change in depth during the acquisition was first performed using the DM system, then the objective was axially moved by the linear stage while keeping the DM in its flat configuration. Both sets of images are shown in Fig. 9, with the first row corresponding to the DM images and second row to the linear stage images. 2P fluorescence images were acquired at a frame rate of 1.1 Hz while illuminating the sample at 1040 nm and collecting all emitted light below 660 nm. Experimental images from Fig. 9 show that the DM system for axially scanning in 2P microscopy provides excellent and comparable results to the ones acquired with the traditional linear stage for the objective.

**Fig. 9.**
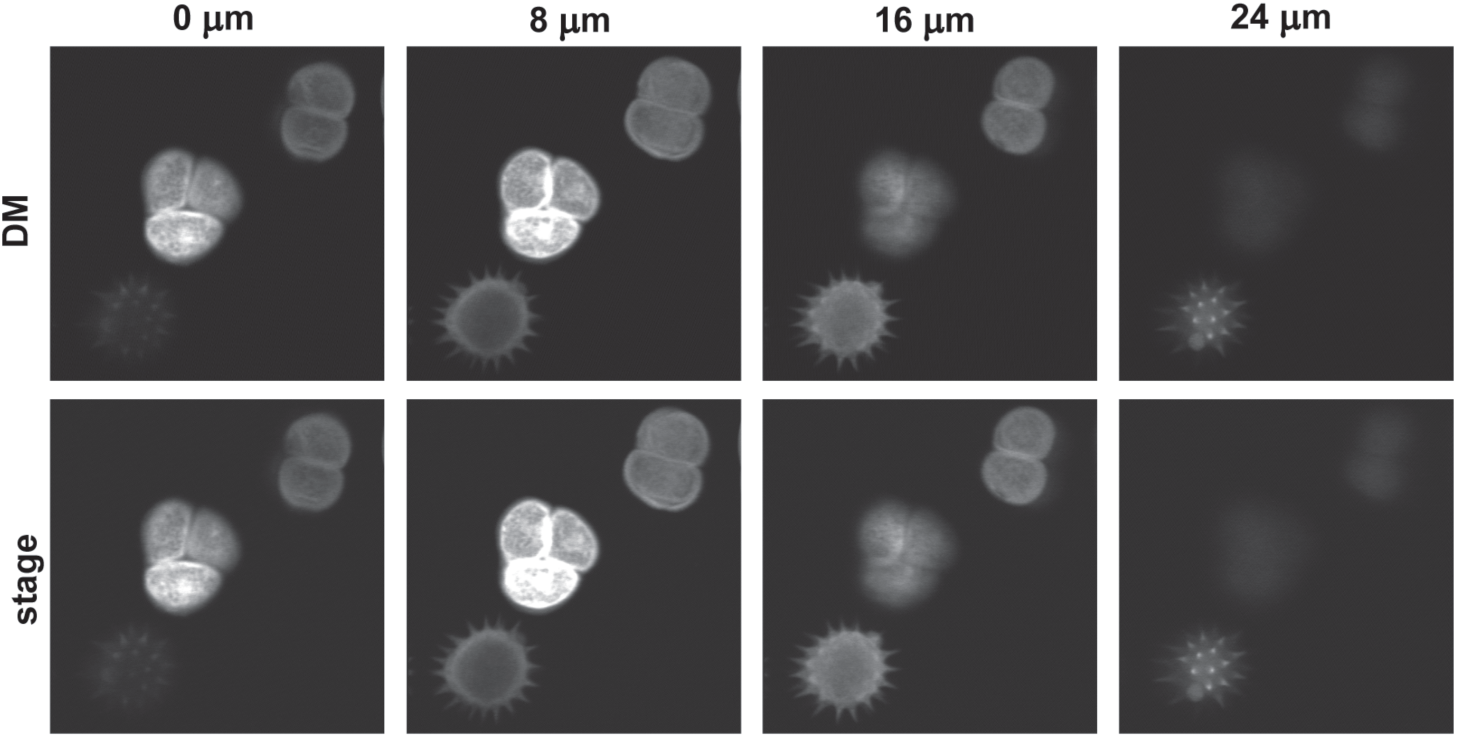
Pollen grains imaged at 1040 nm. 2P images acquired at different *z* planes, using the DM (first row), and the objective linear stage (second row). FOV = 104 µm × 104 µm.

### 4.2. Fixed brain slice

We next imaged a fixed brain slice from an Aldh1l1-tdTomato mouse in which the fluorescent protein tdTomato is expressed solely in astrocytes. The sample was prepared through intracardial perfusion with 4% paraformaldehyde (PFA), which was then sliced coronally at 300 µm. To amplify the fluorescent signal, the tissue samples were then stained via immunohistochemistry and mounted on a glass slide.

Brian slices were illuminated at 1040 nm, and images were recorded at a frame rate of 1.1 Hz. A *z*-stack of 201 images was acquired using the DM, in steps of 0.3 µm. Fig. 10(a) shows a maximum intensity *z*-projection of the recorded *z*-stack, with depth color-coded. There is a complex distribution of astrocyte somas across the whole FOV (288 µm × 288 µm × 54 µm). We show a detailed view of the white box in Fig. 10(a) at 4 given depths, 22 µm, 32 µm, 42 µm and 52 µm, respectively in Fig. 10(b), (c), (d) and (e).

**Fig. 10.**
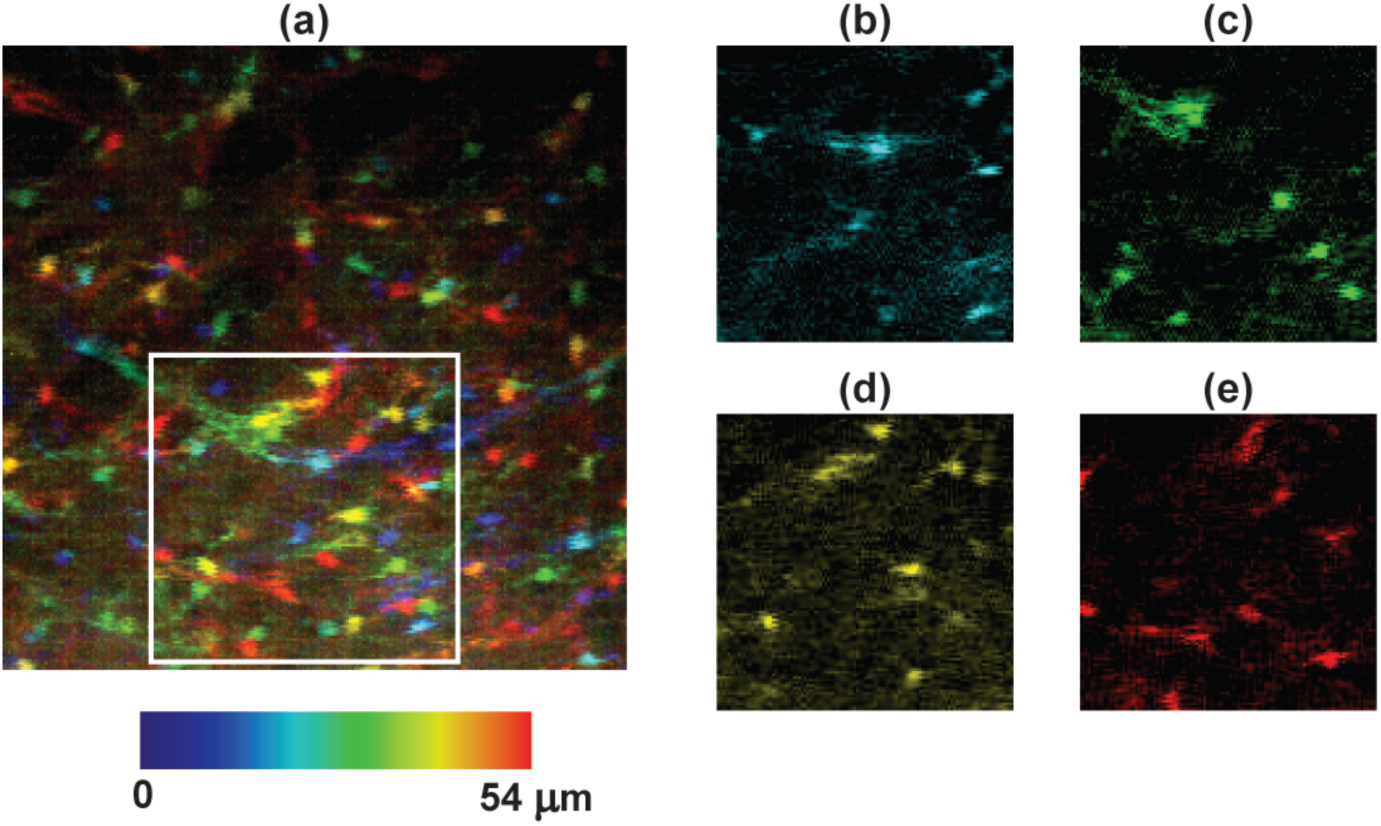
Fixed brain slice (expressing tdTomato in astrocytes) illuminated at 1040 nm. 2P *z*-stack acquired using the DM to change the depth in steps of *δz* = 0.3 µm. (a) Maximum intensity *z*-projection of the *z*-stack, with depth color-coded. FOV = 288 µm × 288 µm × 54 µm. White box is represented for 4 depths: (b) 22 µm, (c) 32 µm, (d) 42 µm and, (e) 52 µm.

### 4.3. In vivo *brain imaging*

Finally, we used the novel 2P microscope for *in vivo* brain imaging. We imaged EAAT2-tdTomato mice to visualize both the soma and the fine processes of astrocytes. On the day of imaging, mice were anesthetized with urethane (100 mg/mL) and a craniotomy (∼3 mm) was made over the imaging area. A 4% agarose solution was placed upon the craniotomy and a glass window was secured into place using dental cement. During the imaging session, mice were kept anesthetized and head-fixed under the microscope objective. We illuminated at 1020 nm, and the image acquisition was taken at 1.1 Hz. A *z*-stack of 201 images was obtained, with steps of 0.3 µm. The ROI was in layer 2/3 of the primary visual cortex.

Fig. 11(a) shows the maximum intensity *z*-projection of the recorded *z*-stack, in which we color-coded depth. As *z* increases, the microscope images in deeper layers. The FOV is 306 µm x 306 µm x 54 µm. In addition, Fig. 11(b), (c) and (d) show the same ROI but at a single specific depth: 8 µm, 30 µm and 52 µm, respectively.

**Fig. 11.**
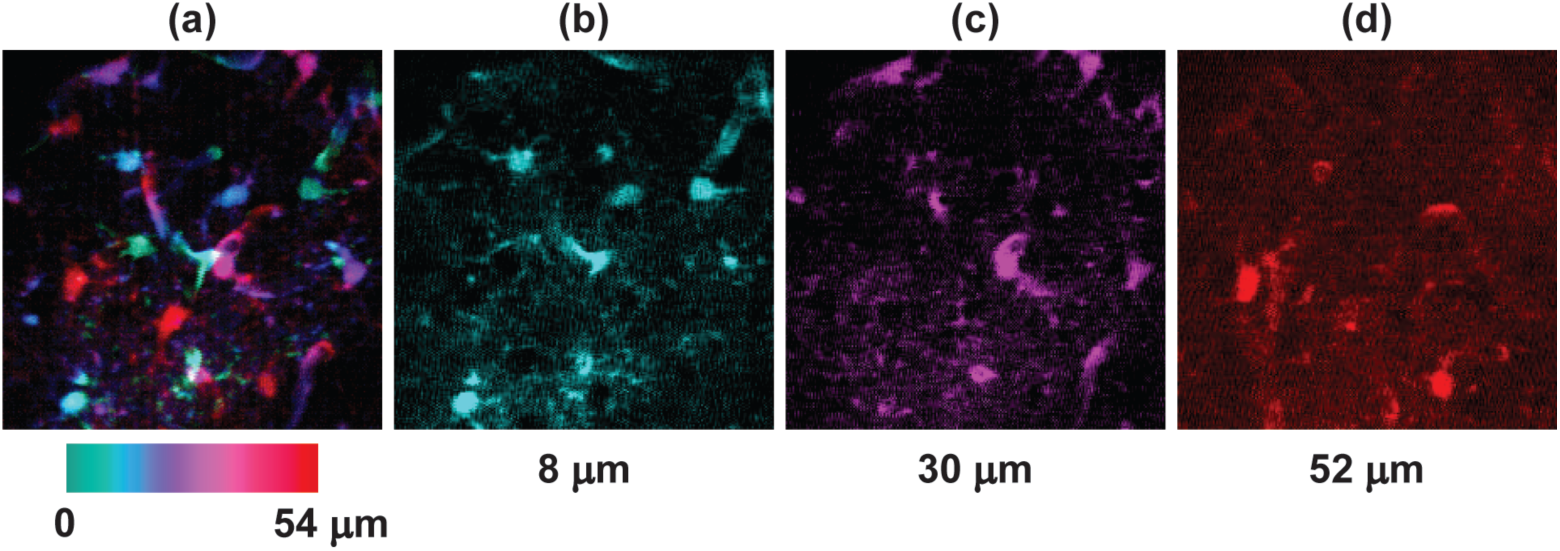
*In vivo* brain imaging of astrocytes illuminated at 1020 nm. 2P *z*-stack acquired using the DM to change the depth in steps of *δz* = 0.3 µm. (a) Maximum intensity *z*-projection of the *z*-stack, color-coded by depth. FOV = 306 µm × 306 µm × 54 µm. Single images taken at depths of (b) 8 µm, (c) 30 µm and, (d) 52 µm.

## 5. Conclusions

This work presents a novel 2P microscope system which incorporates a DM to achieve fast axial scanning. The system is especially advantageous with respect to other volumetric or axial scanning 2P microscopes when imaging more than a few *z*-planes is necessary, and a high axial resolution is required with a reasonable time resolution. In particular, this system is suitable for samples with large populations of cells, as well as for samples containing cells with space-varying signaling such as astrocytes, in which source separation algorithms are not feasible.

By changing the radius of curvature of the defocus implemented on the DM surface, we control the axial position of the beam focus in the sample. The DM used in our system has a fast response time, 100 µs, allowing us to axially scan the sample with a small axial step size without compromising the acquisition frame rate. For example, when combining with a galvanometer mirror scanning the *x* axis at 1.5 KHz, the microscope would scan an *xz* plane with a resolution of 512 x 512 pixels and a dwell time of 1.2 µs, with a frame rate of 2.5 Hz. This could be increased by using a resonant mirror. For example, assuming a resonance frequency of 8 KHz, the microscope could reach frame rates of 7 Hz for the *xz* plane.

We conducted a thorough characterization of the system. First, we experimentally calibrated the PSF of the system for different DM curvatures. Lateral resolution is around 0.9 µm and stays almost constant for all imaging depths, whereas axial resolution is 5–6 µm. As we use the DM with a more negative radius of curvature (*A* closer to −1, for imaging more superficial layers), axial resolution slightly increases because the microscope objective is being underfilled (so the effective NA decreases).

We next calibrated the intensity as DM curvature changes and found that there is a small decrease in intensity reaching the sample as we image deeper layers, due to overfilling the objective. This intensity loss has been calibrated and can be compensated by adjusting the incident beam intensity with the Pockels cell.

The optical setup presented in this paper achieves a total experimental axial scanning range of 54 µm. This range could be increased in the future as DM technology improves and allows to have larger stroke values without compromising the time response. In addition, the user could change the NA of the system. In that way, when reducing the NA, we would increase the axial scanning range, although compromising the spatial resolution. Depending on the sample, the system could be optimized accordingly.

Finally, the instrument was experimentally tested by measuring three different samples. First, pollen grain imaging demonstrated that the DM axial scanning method produces equivalent images as those obtained by axially moving the microscope objective. Next, two neurobiological samples were imaged: fixed mouse brain slices and *in vivo* cerebral cortex of anesthetized mice. Both neurobiological imaging experiments allowed us to visualize and map the morphology of fluorescent astrocytes tangled in depth, showing the capability of this novel technology for axial brain imaging. Future research will focus on the analysis of astrocytic calcium activity along different depths. DM-based 2P microscopy technology opens up the axial dimension for fast brain imaging and it is suitable for temporally and spatially resolving complex and heterogeneous dynamics, including those observed when imaging astrocytic Ca^2+^ sensors, extracellular neurotransmitter sensors, and membrane-targeted voltage sensors in neurons.

## Acknowledgments

Authors thank the members of Poskanzer lab at UCSF for fruitful discussions during the development of this project.

